# Modeling grid fields instead of modeling grid cells

**DOI:** 10.1101/481747

**Authors:** Sophie Rosay, Simon N. Weber, Marcello Mulas

## Abstract

A neuron’s firing correlates are defined as the features of the external world to which its activity is correlated. In many parts of the brain, neurons have quite simple such firing correlates. A striking example are grid cells in the rodent medial entorhinal cortex: their activity correlates with the animal’s position in space, defining ‘grid fields’ arranged with a remarkable periodicity. Here, we show that the organization and evolution of grid fields relate very simply to physical space. To do so, we use an effective model and consider grid fields as point objects (particles) moving around in space under the influence of forces. We are able to reproduce most observations on the geometry of grid patterns. This particle-like behavior is particularly salient in a recent experiment where two separate grid patterns merge. We discuss pattern formation in the light of known results from physics of two-dimensional colloidal systems. Finally, we draw the relationship between our ‘macroscopic’ model for grid fields and existing ‘microscopic’ models of grid cell activity and discuss how a description at the level of grid fields allows to put constraints on the underlying grid cell network.

## 1 Introduction

Grid cells are neurons that activate in correlation with the animal’s position in physical space: they fire only when the animal crosses certain fixed locations in space. For individual grid cells, these firing locations form almost perfect hexagonal grids (Fyhn et al. (2004); Hafting et al. (2005)). The locations of space where a given grid cell is active are called its ‘grid fields’. It was initially thought that grid fields would form such regular patterns throughout the accessible space (Hafting et al. (2005)), regardless of its geometry and of the presence of salient objects such as walls, rewards or threats. However, recent experiments refuted this view of a *perfect and universal grid*. When recorded in very large environments, grid patterns display non-uniformities (Stensola et al. (2015)) and walls can influence grid orientation and distort it from a perfect hexagonal lattice (Stensola et al. (2015); Krupic et al. (2015); Carpenter et al. (2015)).

Also the presence of goals can modify grid patterns (Boccara et al. (2016)). Finally, grid patterns evolve under manipulations of the environment, like stretching or squeezing (Barry et al. (2007)). A recent experiment explored this adaptability of grid patterns. Given that walls can fragment grids into separate pieces that do not form a coherent grid, Wernle et al. (2018), asked what would happen if a separation wall between two such incoherent pieces is removed. They observed that, rather than forming a totally new grid, existing grid fields shift and adjust, as if finding a compromise between coherence and conservation of the patterns. Grid fields located closer to the former separation wall ‘move’ more to establish coherence, whereas grid fields far away from the separation wall remain located. This experiment provides crucial new cues about grid field dynamics.

So far, explaining grid cell properties has been attempted through models of neural networks. These models aim at deriving a causal relationship between certain properties of neurons and the observed grid patterns. Here, we refer to them as *‘microscopic models’*, because they describe the emergence of grid patterns at the microscopic level (neurons, synapses, firing dynamics, plasticity,…). These models advanced our understanding of grid cells. Yet, so far no model has succeeded in accounting for all observed phenomena—in particular the effects of boundaries and geometry on grid patterns are scarcely described.

Here, we take a different approach, forget for a time about the neurons, and directly consider grid fields as point objects (particles). We assume these particles to move smoothly in space under the influence of forces, friction and noise. This picture naturally comes to mind when looking at the time evolution of grid patterns in experiments— particular in the merging experiment by Wernle *et al*. Hence, we build a model on this impression to see how far it can represent what we see. We call this representation a *‘macroscopic model’*, since it takes place at the level of macroscopic observables (grid fields, physical space) rather than microscopic elements (neurons etc.). Because of the presence of noise and friction, we are in the framework of colloidal systems in two dimensions—a field of physics where many results are already known. This provides us with conceptual tools to understand and interpret our results.

This approach requires to find an expression for the motion of particles and the forces that act on them. This will be the object of Section 2. In Section 3, we make clarifications on the notion of grid symmetry and its measure. Then, we present the outcome of numerical simulations in setups that either reproduce experiments conducted on rodents (Section 4) or predict the outcome of experiments not performed yet (Section 5).

In Section 6, we relate the macroscopic description to existing microscopic models. Finally, we discuss the relevance of a macroscopic description.

## 2 The model

### 2.1 Grid fields from one grid cell

Let us first consider grid fields from a single grid cell. In our model, its grid fields are seen as a system of *N* particles that interact with each other and with the environmental borders, with noise and friction, as illustrated in Fig. 1. The underlying intuition is that the fields behave as colloidal particles moving around in a substrate (Manoharan (2015)).

**Figure 1.**
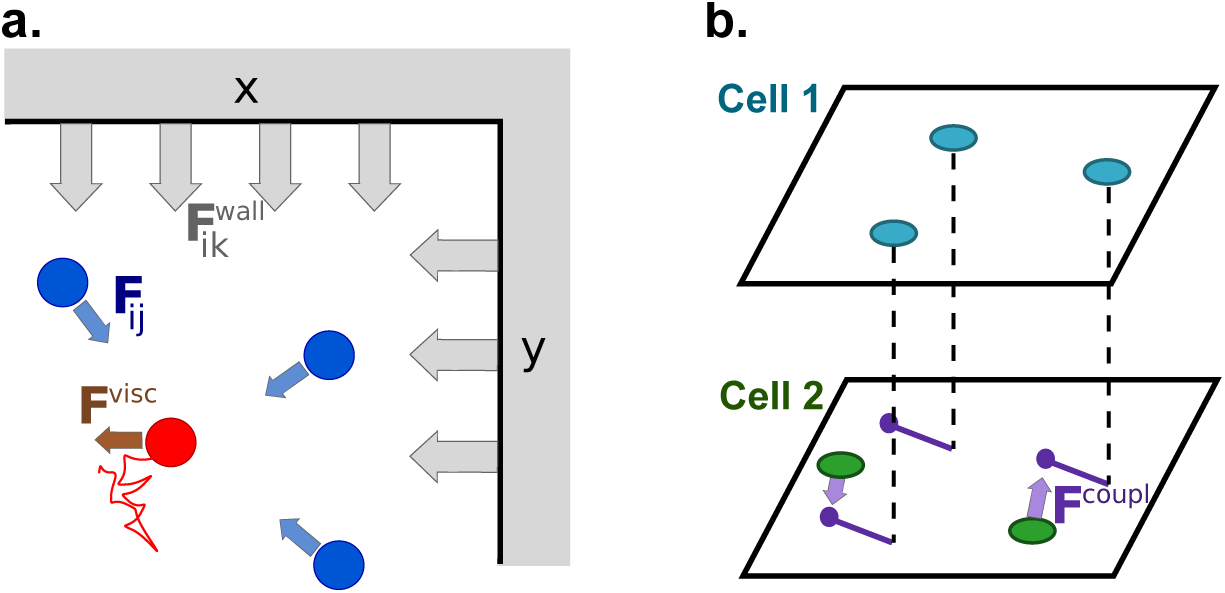
Schematic representation of the model. **a**: grid fields from a given cell. A grid field (red) moves under the influence of forces coming from other grid fields (blue), walls (gray), viscosity (brown) and noise (related to the viscosity through the Einstein relation). **b**: coupling between two cells from the same module is introduced as an attractive force from the fields of one cell (cell 1, blue fields) on the fields of another cell (cell 2, green fields) at a certain offset position from field of cell 1 (purple)

We describe the system by a Hamiltonian that governs the time evolution of particles at temperature (i.e. noise level) *T*:

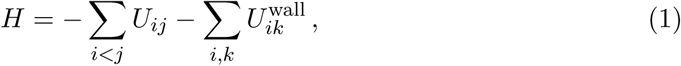

where *U*_*ij*_ is the potential between particle *j* and *i*, and 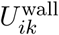 is the potential between wall *k* and particle *i*. The corresponding Langevin equation, for particle *i* is

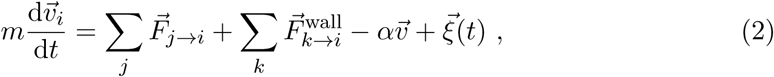

where the forces derive from the potentials above and 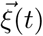 denotes Gaussian noise of variance 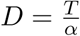 (Einstein relation at temperature *T*).

We use forces with a Gaussian kernel, as detailed in the Supplementary Material. The forces between particles are repulsive. As for the forces originating from the walls, we will demonstrate that an attractive force describes the experimental observations better than a repulsive force.

### 2.2 Grid fields from several cells from the same module

The model above only considers grid fields from a single cell, regardless of other grid cells. However, it has been experimentally demonstrated that grid cells are grouped into ‘modules’, which are ensembles of cells organized along the dorso-ventral axis of the entorhinal cortex (Stensola et al. (2012)). Grid cells from the same module form patterns of the same spacing, the same orientation and random phases. Between modules, phase offsets and orientations are seemingly uncorrelated while spacings increase dorso-ventrally.

We reasoned that this coupling between cells can relate with the suggested single-cell coupling between fields in two ways: either the effective force between fields is a network effect and therefore already contains the coupling between cells, or the effective force between fields comes from another mechanism and the coupling between cells adds on top of it. To implement the possibility of a varying degree of independence between grid cells, we consider two grid cells and their grid fields. The dynamics of the grid fields of the first grid cell follow the single-cell equation (2). The grid fields of the second grid cell follow the same dynamics with an additive term of coupling:

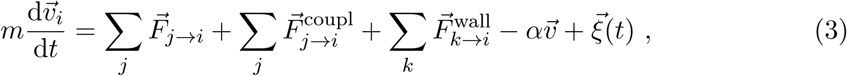

where 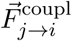 is the force exerted by particle *j* from cell 1 on particle *i* from cell 2. We expect that for high coupling forces, the patterns of cell 1 and cell 2 will be perfectly aligned, whereas for weak coupling forces, the patterns will be independent.

We choose to take 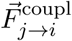 as an attractive force towards positions that correspond to the fields of cell 1, translated by an arbitrarily chosen offset (Figure 1b).

Simulation details and parameter choices are reported in the Supplementary Material.

## 3 Preliminary remarks on grid symmetry and gridness scores

### 3.1 The classical gridness score

Before going further, we have to specify how grid symmetry is defined.

In experiments, a given neuron is considered as a grid cell if its firing pattern has a certain degree of hexagonal symmetry. Since the discovery of grid cells and their hexagonality, hexagonality is usually quantified by a measure called *‘gridness score’* introduced by Sargolini et al. (2006) and subsequently slightly modified by Langston et al. (2010). A cell’s gridness score is computed from its smoothed rate map, by taking this rate map’s autocorrelogram, extracting a sub-ring of it and comparing this sub-ring to rotations against itself. The details of this definition are given in the Supplementary Material. A given cell is then considered a grid cell if its gridness score exceeds a threshold, most commonly either 0 or the 95th percentile of the distribution of gridness scores from a shuffled dataset.

Gridness scores have been extensively used both to define grid cells and to evaluate their quality. Here are a few remarks on the information they provide:

1. By definition, the gridness score is a global, and not local, measure of hexagonal symmetry.
2. Although from their mathematical definition gridness scores could range between 2 and +2, in practice a perfect grid cannot exceed ≈ 1.5. We estimated this upper bound by generating ideal grid maps (putting bumps of activity on a perfect hexagonal grid—no model involved) and computing their gridness scores (Fig. 2a). This upper bound is due to the choice of the radius of the inner disk that leaves a region of the autocorrelogram always correlated with its rotation, which blurs the signal (we have checked that removing this region restores the upper bound to 2).
3. Experimental gridness scores do reach this upper limit, quite remarkably. See Fig.1 in (Sargolini et al. (2006)).
4. Does a high gridness score necessarily indicate a strong hexagonal symmetry? To answer this question we generated artificial rate maps of different shapes and symmetries.
  - Mere equidistance between fields (without 6-fold order) is not enough to account for a high gridness score (Fig. 2b).
  - However, starting from perfectly hexagonal patterns and perturbing them randomly, we observe that gridness scores are quite robust to such distortions. In particular, if these deformations are applied on the edges, gridness scores are almost unchanged, indicating that the symmetry captured concerns mostly the center of the box.
  - In some cases where we perturb the edges, the distorted grid has a higher gridness score than the perfect grid (Fig. 2c). The explanation is related to point 2: what prevents gridness scores to be high is the inner ring of the autocorrelogram that has a constant value. Some limited jitter on the edges lowers the correlation of this area with rotations of itself while not altering the anticorrelation of the outer ring with rotations against itself, hence increasing the gridness score. Such a counter-intuitive effect reflects the complexity and indirectness of the measure.
  - In conclusion, decent gridness scores (> 1) can come from quite bad-looking patterns. Yet the highest gridness scores (> 1.4) could only be accounted for by grids that are near to perfect, at least in the center of the box (Fig. 3).
  - The reported gridness scores are measured in relatively small environments (boxes of less than 2m side length). Therefore the level of symmetry they reflect regards *local* order. In larger environments, global gridness scores have not been reported. Yet, even in theses small areas, global order is broken (see Fig.4i in (Stensola et al. (2015))). We will come back to this point in Section 4.1.
  - The autocorrelogram is the standard way to characterize grid patterns, not only to compute gridness scores, but also to define grid spacing, orientation, ellipticity, ‘shearing’ (Stensola et al. (2015)), etc. However, autocorrelograms can cancel out distortions at the border of the box. We will come back to this point in Section 4.3.
  - Experimental gridness scores are distributed along all possible values. See for instance Supplementary Fig. 5 in (Bonnevie et al. (2013)). Yet, from a computational point of view, existing grid cell models have trouble accounting for all the points cited above: either they produce only ‘super grids’ of very high gridness score, or they produce a wide range of gridness scores but not up to the maximum.

**Figure 2.**
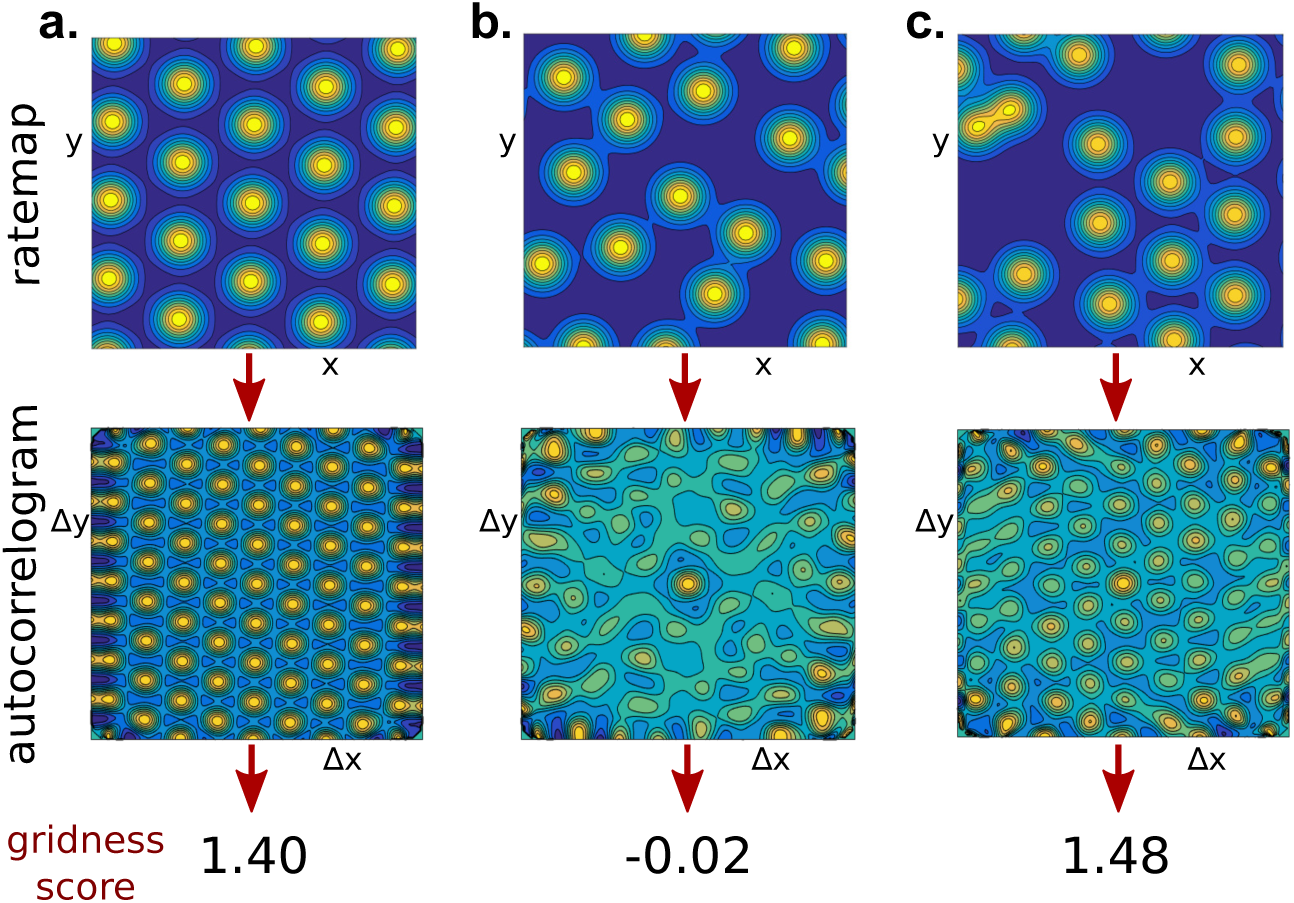
Illustration of some features of the gridness score. Examples of generated rate maps (top), their autocorrelograms and gridness scores (bottom). **a**. Perfect hexagonal patterns lead to gridness scores saturating around ≈ 1.5. **b**. Bumps of activity scattered randomly at roughly equidistant positions lead to very low gridness scores. **c**. However, very high gridness scores can be obtained by perturbing an initially perfect grid. The perturbed grid sometimes even has a higher gridness score than the unperturbed grid. See Fig. 3 for a systematic exploration of this effect

**Figure 3.**
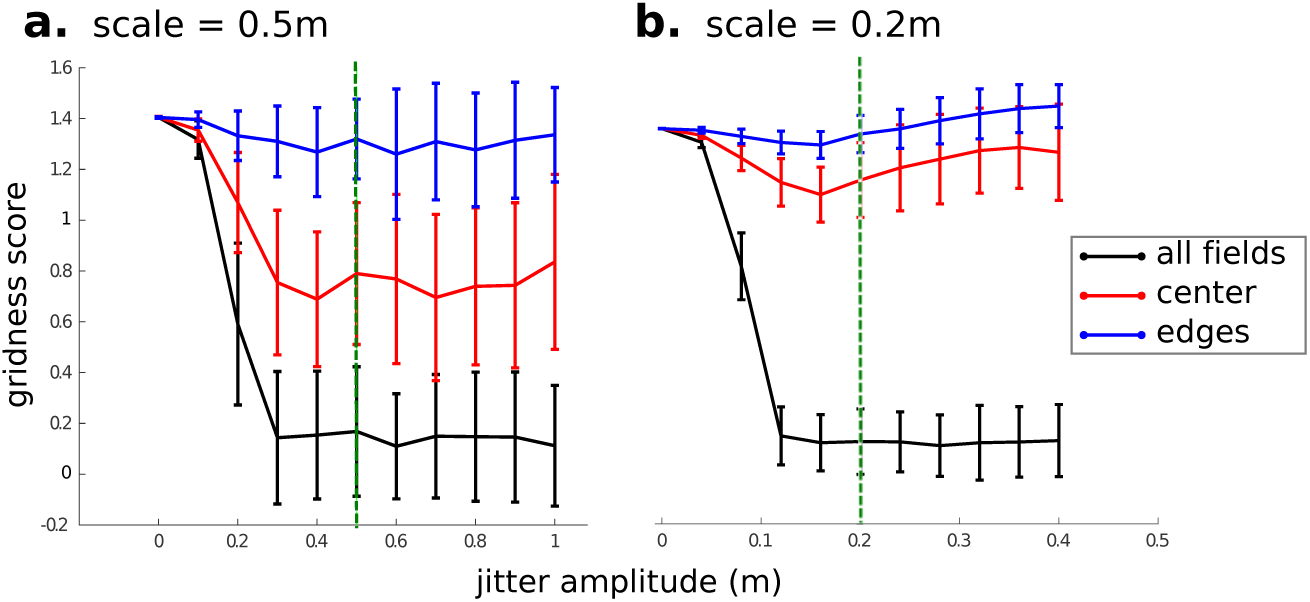
Gridness scores are robust to perturbations around perfect hexagonal grids. We generated perfect hexagonal grids of random orientations and of spacing *L* = 0.5m (**a**) and *L* = 0.2m (**b**), in a 2m × 2m box. We then jittered the fields randomly within a disk of varying radius (*x*-axis). For each jitter amplitude, we computed the resulting gridness scores over 100 realizations (*y*-axis, error bars indicate standard deviation). Green dashed lines indicate grid spacings. Black: all grid fields were jittered. Red: only grid fields in the center of the box were jittered (at distance > 0.3m from the walls, i.e., half of the total area.). Blue: only grid fields on the edges were jittered (at distance < 0.3m from the walls). Note that gridness scores remain particularly high in the latter case, and can even *increase*, for very strong perturbations. See text for an explanation

**Figure 5.**
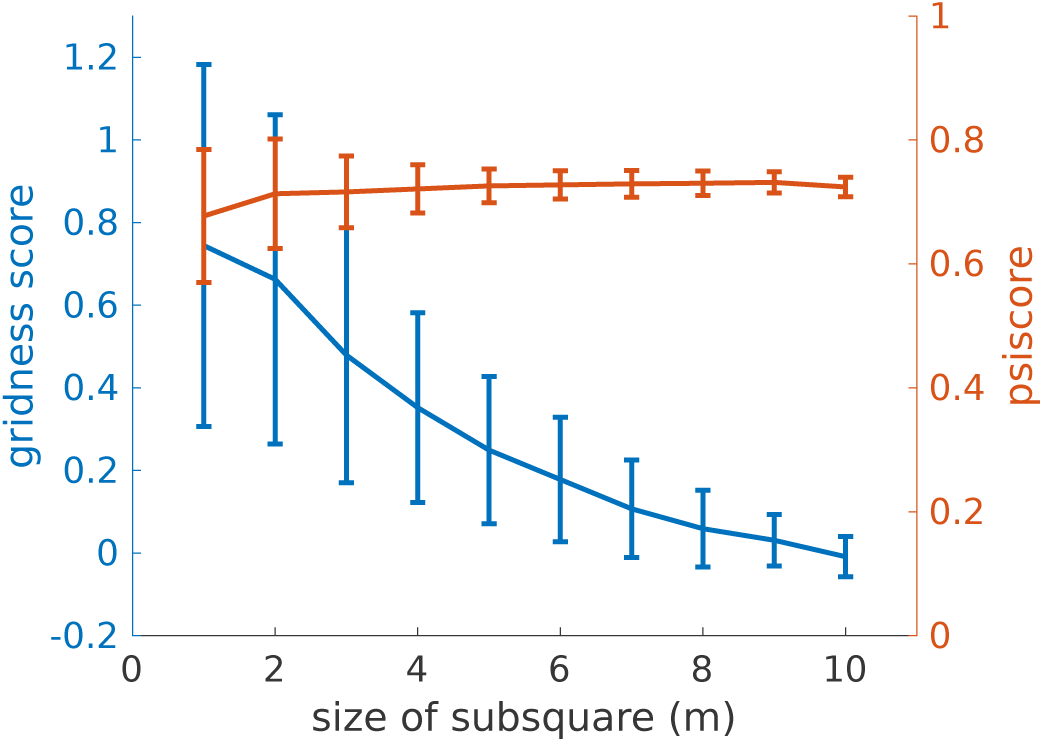
Even with large numbers of fields, hexagonal order is only local in our model. Simulations were run on a 10m × 10m torus as in Fig. 4b. Gridness scores (blue) and *Ψ* scores (orange) were computed on a sub-square of increasing size. The gridness score, which is a global measure, drops when increasing the size of the box, while the average *Ψ* score, a local measure, does not depend on the size of the box. Error bars indicate standard deviations over 100 simulations

### 3.2 An alternative measure

Put together, these facts point to the utility of defining a local measure for grid symmetry, that captures local distortions near the walls and is robust to long-range order breaking in large environments (Stensola et al. (2015)). It turns out that such a measure has already been introduced in the context of the physics of 2-dimensional systems (Halperin and Nelson (1978)). It is called the orientational order and writes locally

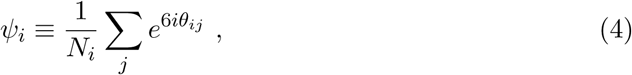

where *N*_*i*_ is the number of nearest neighbors of unit *i* and *θ*_*ij*_ is the angle of the vector linking *i* to its neighbor *j*.

The absolute value of the complex number *Ψ*_*i*_ quantifies how much *i*’s neighbors arrange hexagonally. The argument of *Ψ*_*i*_ gives the *local orientation* of the grid. So far grid orientation has always been measured globally from the autocorrelogram, ignoring the possibility of a non homogeneous orientation^1^. Weber and Sprekeler (2018b) have recently applied this measure to spike maps from grid cells and showed local grid distortions and changes in local grid orientation (code available at (Weber (2018))).

At the level of a whole pattern, the orientational order is the average of the local *Ψ*_*i*_:

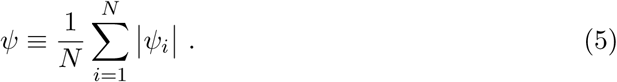

Having clarified the notion of grid symmetry, we can now see how our grid field model accounts for what is observed in experiments.

## 4 Simulation results - verifications of experimental facts

### 4.1 Formation of grid patterns

The main characteristic of grid fields is that they arrange in a hexagonal pattern. We first outline the conditions under which they do so in the single-cell version of our model (Eq.1). We make the following observations:

- **For moderate numbers of particles** (*N ≾* 30, i.e. the number of grid fields typically observed in a standard experimental recording) we only observe grid-like patterns if we assume *repulsive* walls. Indeed, with the assumed repulsive force between particles, something has to ‘contain’ them in the box. When we impose repulsive walls, the grid fields arrange with almost regular spacing and locally form hexagons (Figure 4a). However, the grid scores are quite low in comparison to experiments (values go up to 1.5, see for instance Refs. Sargolini et al. (2006); Bonnevie et al. (2013)).
- To obtain higher grid scores, we use **large numbers of particles.** To focus on pattern formation and not on edge effects, we first simulate the system on a large square torus (10m × 10m, Fig. 4b) and study wall effects in the next paragraph. We look at *local* grid order on a subset of the system corresponding to experimental length scales. Here, the gridness score is computed on a sub-square of 1m × 1m. This local gridness score is high (Fig. 4b). We conclude that hexagonal pattern formation in our model is a large-*N* effect. In the rest of this paper, when simulating grid fields in a box, we will thus always assume that they are surrounded by a large number of fields outside of the walls.
- The observed grid order in our model is local. Global order is broken (Fig. 5). From a neuroscience point of you, this is strongly reminiscent of the breaking of global grid order observed in large boxes (Stensola et al. (2015)). From a physics point of view, it is known that there is no long-range positional order in two dimensional crystals (Peierls (1935); Berezinskii (1971); Kosterlitz and Thouless (1973)). Our results suggest that the latter may explain the former.
- If we increase the temperature *T*, the system enters a disordered phase. To obtain hexagonal patterns, we will thus focus on low temperatures.

**Figure 4.**
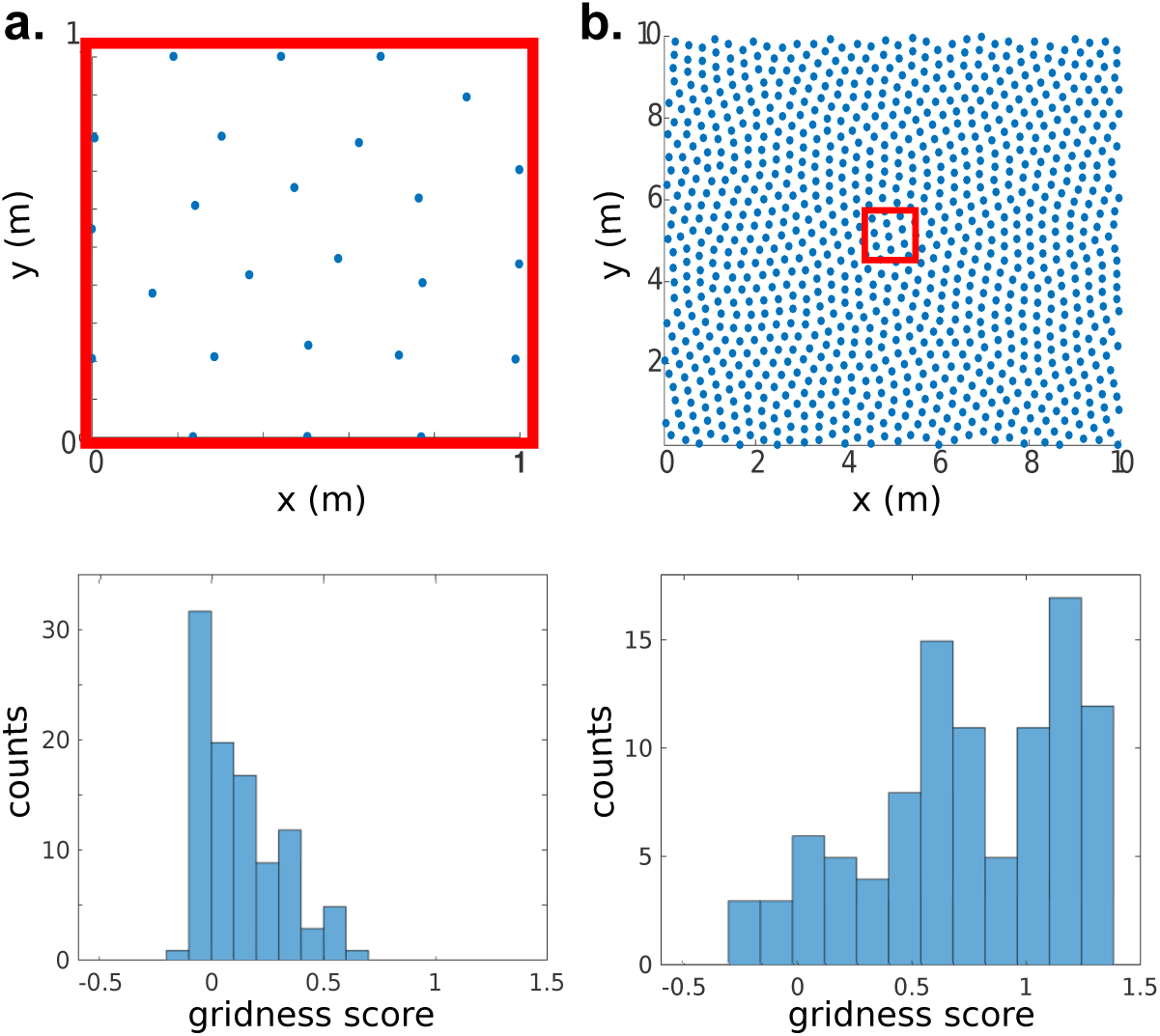
In simulations of our model, hexagonal grid order requires a large number of particles. Top row: examples of patterns in the asymptotic state. The area in which the gridness score is computed is indicated with a red square. Bottom row: corresponding gridness score distributions over 100 simulations. **a.** A moderate number of grid fields (in a 1m × 1m box) leads to evenly distributed patterns of poor hexagonal symmetry (low gridness scores). **b.** A large number of grid fields (in a 10m × 10m torus) leads to strong local order. The gridness scores measured in a sub-box of 1 × 1m (red square) reach high values if the area does not contain breaks in long-range order, and low values if it does

### 4.2 Module coherence

#### 4.2.1 Simulation of the multiple cell model

We now present simulations of the model with two cells (Sec. 2.2) with a varying intensity of the coupling force 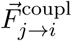. At the end of each simulation we measured the coherence (or rather the incoherence) *I*_mod_ between patterns and averaged over many simulations (Figure 6). *I*_mod_ is defined as the minimal average distance between one pattern and any spatial translation of the other, see Supplementary Material for details.

**Figure 6.**
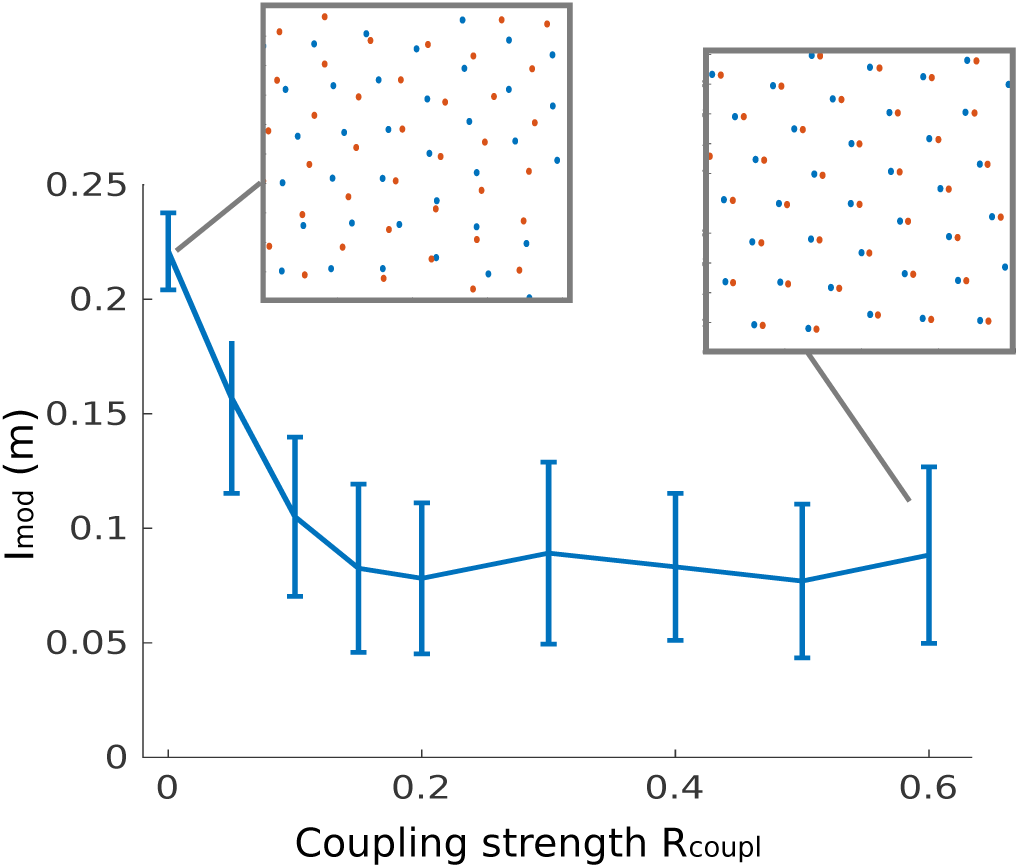
Incoherence *I*_mod_ between patterns as a function of the coupling strength between two cells on a 10m × 10m torus. Incoherence is measured as the minimal average distance between one pattern and any spatial translation of the other and 0, if the two patterns are a translation of each other, see eq. (8) in Supplementary Material. We can see that a saturation is reached around a strength *R*_coupl_ = 0.1*R*_rep_ (see the full expression of the forces in the Supplementary Material). Error bars indicate the standard deviation across 100 simulations. Insets show examples of the obtained patterns (zoomed on a 2 ×2m sub-square) for *R*_coupl_ = 0 and *R*_coupl_ = 0.6 respectively

We observe that a weak coupling (1/10 of the strength of the repulsion between fields) suffices to align patterns.

Interestingly, almost perfectly aligned patterns emerge even for distorted grids, i.e, hexagonal symmetry and module coherence decouple. The presence of both coherent and distorted grids is an experimentally testable prediction of our model.

In experiments, grid patterns recorded from the same module have a degree of coherence much higher than expected by chance (Stensola et al. (2012)). But small deviations from a rigid offset occur, up to a limit that remains to be quantified. The measure *I*_mod_ we introduce here could be applied to experimental maps as an estimate of how far they depart from perfectly coherent patterns, and draw a closer relationship with our model.

#### 4.2.2 Simulation of repulsing hexagons

In our effective multiple-cell model with attractive forces between grid fields of different cells we showed that grid orientations align, as observed in experiments. Yet, in this case the phase offset between grids is fixed and thus this model cannot reproduce the observation that the phases of different cells in the same module are homogeneously distributed (Hafting et al. (2005); Yoon et al. (2013)).

We now take a different approach in which the fields of different grid cells do not attract each other, but repulse each other. To simulate more than just two cells (with reasonable simulation times) and to decouple the emergence of hexagonal patterns and grid-alignment, we ask how rigid hexagonal patterns would organize if put in repulsive interaction.

To this end, we treat recurrent inhibition between grid cells as an optimization problem. We consider a set of perfect grid cells with identical grid spacing. Each ‘cell’ is a hexagonal arrangement of Gaussian fields. The orientation and phase of one cell is fixed. The orientations and phases of the remaining cells are parameters that are determined such that the overlap of the firing fields of all grid cells is minimal. Details on the simulations are provided in the Supplementary Material. We observe that the overlap is minimized if all grids have the same orientation, but different phases, as shown in Fig. 7.

**Figure 7.**
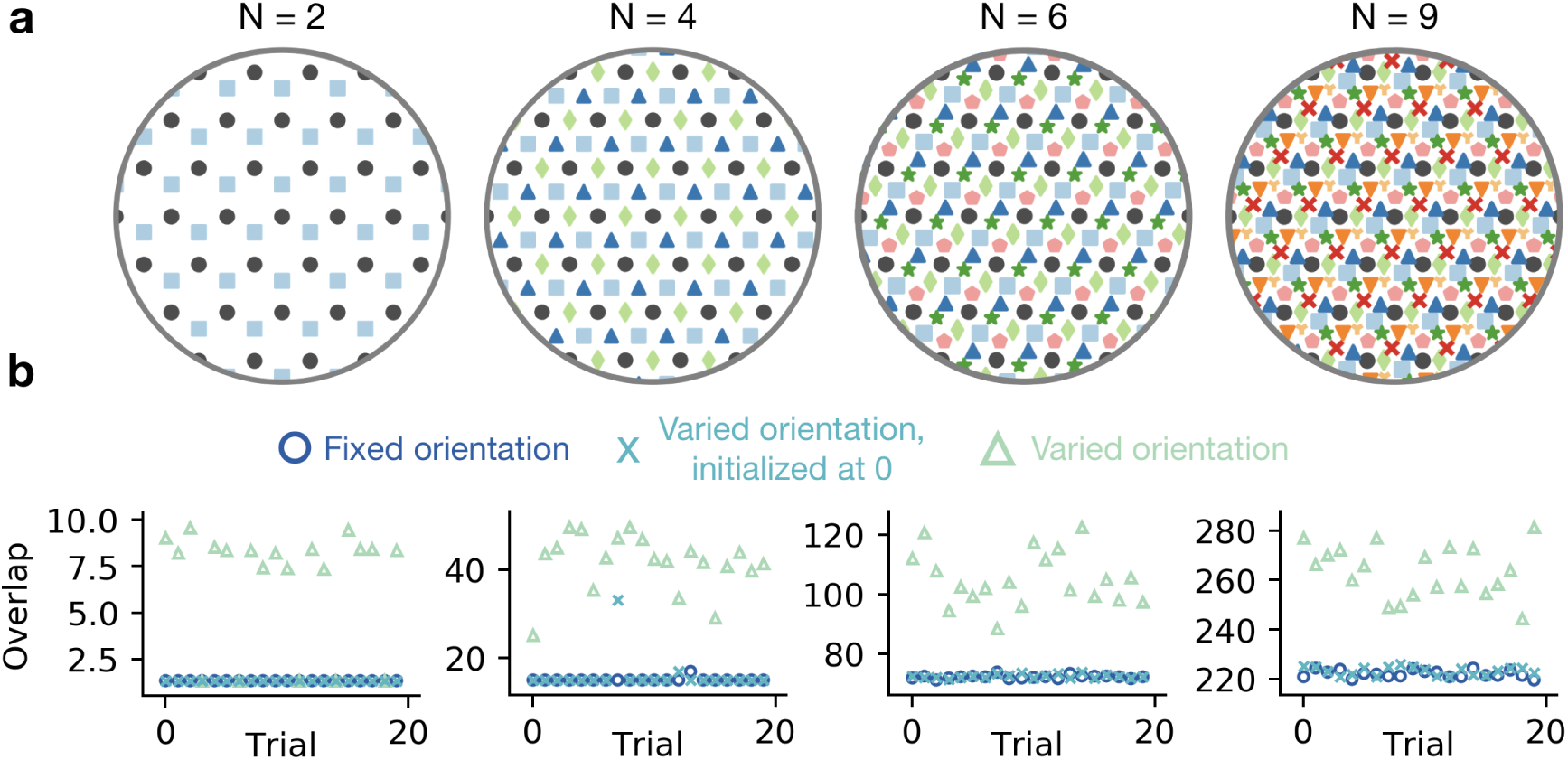
Co-orientation and phase spreading in repulsing grids: preliminary results in simulations where perfect grids try to minimize the overlap between their grid fields. (**a**) Center locations (colored symbols) of *N* perfectly hexagonal grids of Gaussian fields after the overlap between all grids has been minimized numerically. Dark gray dots correspond to the reference grid, which is static. (**b**) Overlap (in arbitrary units) of Gaussian fields for 3 × 20 minimization trials for the different values of *N* shown in **a**. In one set of 20 trials (dark blue circles), all grid orientations are fixed at the same value and only the grid phases are optimized in the sense of minimal overlap between grid fields. In a second set of 20 trials (medium blue crosses), the minimization algorithm varies not only the phase but also the orientation of the grids and all orientations are initialized at 0 degrees. In a third set of 20 trials (light blue triangles), again all phases and orientations are varied, but here orientations are initialized at random. For all *N* considered, the solution with minimal overlap corresponds to a scenario of aligned grids (all orientations 0). In all trials, the phases are initialized at random

These preliminary results support the idea that the observed grid co-orientation and phase spreading arise from mutual inhibition between grid cells of the same module. Further work will have to quantify how much this inter-cell effect interacts with pattern formation in individual cells.

### 4.3 Effect of the walls

What kind of force, if any, do the walls exert on grid fields? In Section 4.1 we have shown that walls are not required to ‘contain’ the grid fields, since grid patterns could also emerge in our model from the interaction with a large number of grid fields outside of the box. Therefore, walls would only perturb the already existing (and already distorted) grids.

Experiments revealed that walls have an influence on both grid geometry (grids are less hexagonal, more elliptic, or sometimes even break up) and grid orientation (grids tend to cluster around an angle of roughly 8 degrees relative to the nearest wall) (Stensola et al. (2015); Krupic et al. (2015)).

Stensola et al. (2015) proposed that both effects originate from the same cause, namely a ‘shearing force’ exerted by the walls on an originally hexagonal grid pattern. Note that this proposed explanation is given at the macroscopic level: it describes how walls effectively affect grid fields.

We decided not to use a shearing force in our model, because it would mean fine-tuning the shearing parameters in order to reproduce experimental features measured on the autocorrelogram. Also, shearing forces do not have clear mechanisms at the cellular level.

Instead, we assume that walls exert an *orthogonal attraction* on the grid fields. This is motivated by two independent arguments:

- From a bottom-up point of view, we know that there are excitatory border cells that fire close to walls (Solstad et al. (2008)). Intuitively, the most straightforward guess is that these cells have a tendency to excite grid cells to fire closer to the wall, i.e., to exert an orthogonal attraction on the grid fields. This remains to be experimentally demonstrated, though.
- From a top-down point of view, we observed that attractive walls induce what Hägglund (2017) have called ‘barrel distortions’ of the grids in our simulations: a dilatation of the grid in the center, accompanied by a squeezing on the edges and a bending of the axis (Fig. 8).

**Figure 8.**
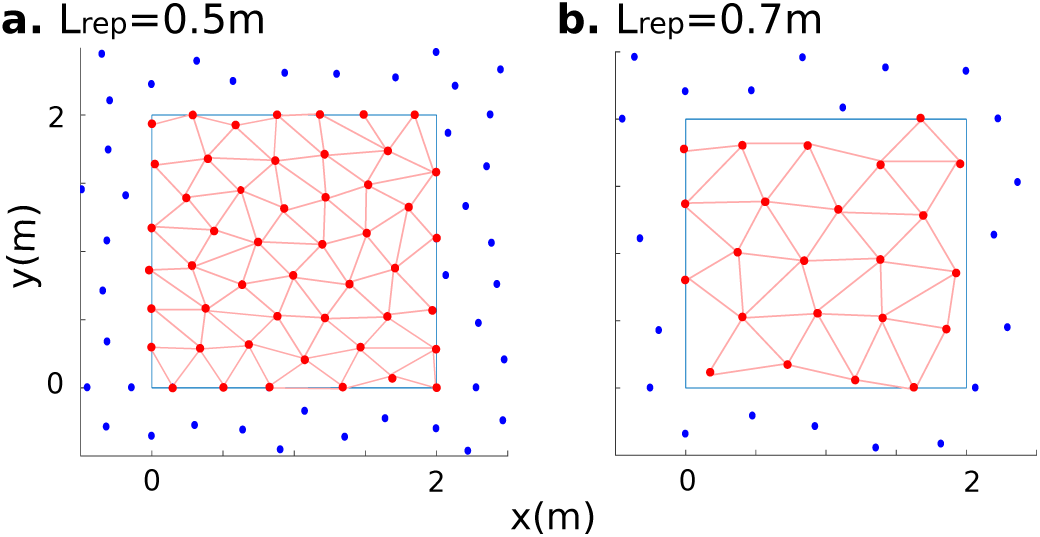
Examples of the effect of attractive walls on grid patterns, reminiscent of the distortions observed in experimental grid patterns, for ranges *L*_rep_ = 0.5m (**a.**) and *L*_rep_ = 0.7m (**b.**) in a 2m × 2m sub-box surrounded by a large number of fields (see Supplementary Material for the full expression of the forces). For clarity we represent only a fraction of the total 10m × 10m torus. Blue: some of the fields outside the box. Red: fields inside the box. Pink lines serve as a guide to the eye. Note the bending grid axis and the tendency of grid fields to stick to the walls, both due to the attractive force coming from the walls

The effect of walls on grid patterns still remain to be better characterized in experimental data to test our assumption of attractive walls. The *Ψ* score could be of help for this purpose.

### 4.4 Response to wall removal

We now turn to the wall-removal paradigm described above.

Wernle et al. (2018) showed that, before the wall is removed, the initial patterns form two grids of roughly the same spacing and orientation^2^ but different offset: they are thus called ‘incoherent grids’, because they do not form a single hexagonal pattern. After wall removal, the grid fields ‘merge’: they rearrange in the center of the box but remain fixed at the edges. This persistence of the initial pattern in spite of its changes in other parts of the box has been described as an ‘anchoring force’ ^3^.

How can we model this anchoring force? The attraction from the walls that we used so far would not suffice because it is not localized enough. Here we take another approach, inspired by our previous observation that grid fields outside the box also have to be modeled. We make the following assumption (illustrated in Fig. 9): everything evolves as if, before wall removal, grid fields inside the A (resp. B) compartment ‘saw’ fields outside the box that are arranged on a perfect grid with offset *𝒪*_*A*_ (resp. *𝒪*_*B*_). After wall removal, fields rearrange inside the box but still see the same fields outside the box. Indeed, this is a very simple way to model the anchoring of grids, consistent with the idea that anchoring probably comes from distal cues. Of course, this approach is one among many. In particular, it neglects many aspects (wall effects, the irregularity of the grid, the occasional merging of two nearby fields, etc.) for the sake of simplicity.

**Figure 9.**
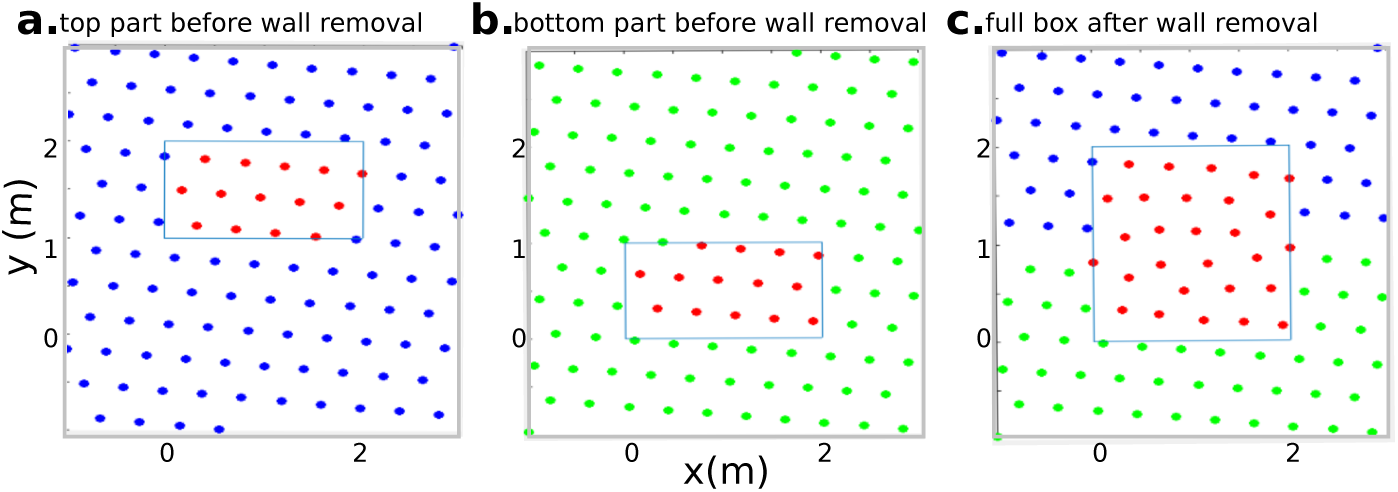
Implementation of the wall-removal setup. Before removing the wall (a-b), the fields in compartment A find their positions consistently with a grid of a certain orientation, spacing and offset (a); while the fields in compartment B do the same with a grid of same orientation and spacing but different offset (b). After removing the wall, the fields from both compartments ‘see’ each other, while they keep the anchoring of the former grids across the remaining walls (c)

Fig. 10 shows how the patterns evolve after removing the wall: grid fields in the central part of the box move more (Fig. 10b) than grid fields close to the boundaries (Fig. 10a). This evolution of fields is such that a local grid order is found (Fig. 10c). Qualitatively, our simulations lead to similar results as observed experimentally. Tn terms of hexagonality (measured indirectly as the standard deviation of neighbor distances; Fig. 10c), we get more regular grids because of our simplifying assumptions that leave aside many possible grid distortions. However, in terms of pattern correlation and normalized field displacement (Fig. 10a and b), the qualitative agreement is good, without fine tuning of parameters (we only choose a grid spacing comparable to experiments).

**Figure 10.**
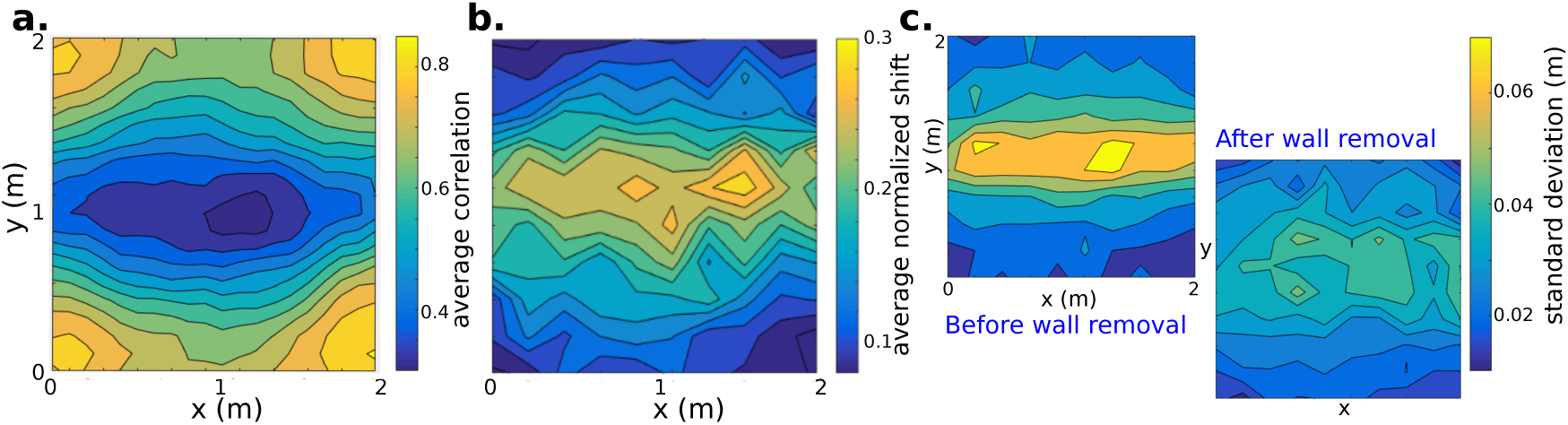
The model in the merging setup, averaged over 100 simulations. **a**: average correlation of the grid pattern after wall removal with the one before wall removal, showing that the grid fields rearrange in the center of the box while being ‘anchored’ on the borders far from the former separation wall. **b**: average displacement of each grid field as a function of its initial position, normalized by the grid scale. **c**: The motion of grid fields is such that grid coherence increases near the separation wall. Standard deviation of the distances of each grid field’s neighbors as a function of its position, both before and after wall removal

In conclusion, the experimental observations can be accounted for very simply. Importantly, note that in this particular case we do not even need interaction with the walls (we can neglect the grid deformation) nor additive forces on the fields: the simple presence of the grid outside the box is sufficient to keep the memory of the grid inside. This is reminiscent of the hypothesized role of distal cues as a reference in grid alignment. These fixed outside grids are an effective—and efficient—way to model anchoring.

## 5 Simulation results - predictions

### 5.1 Local and global grid order

Our results from paragraph 4.1 allow us to derive several predictions regarding experimental grid patterns.

First, in very large environments, global order is broken, therefore the gridness score measured in a subpart should be a decreasing function of the size of this subpart (Fig. 5).

Second, the breaking of global grid symmetry is a consequence of its two-dimensionality; it is not an effect of the walls^4^. Therefore, if our model is correct, all the results on global order breaking in large environments should be independent of the geometry of the box. For instance, a large circular enclosure and a large square enclosure should lead to the same decrease of the gridness score as a function of the size of the portion on which it is measured.

Third, the local order should be high. This is easily verified by applying local orientational order *Ψ* to experimental grid maps (Weber and Sprekeler (2018b)).

### 5.2 Module coherence

Our model predicts that hexagonal order and module coherence are two independent features. Therefore, one should find in experiments:

- examples of quite regular grids from the same module that are not well aligned to each other (in particular near the walls, whose force can overcome the alignment force),
- examples of poorly hexagonal grids from the same module that nevertheless are well aligned.

This prediction requires to be able to quantify both the local hexagonality and the grid alignment: this could be done by the two measures *Ψ* (see also Weber and Sprekeler (2018b)) and *I*_mod_, which we have propose above.

### 5.3 Density of grid fields

In paragraph 4.3, we assumed that walls exert an attractive force on grid patterns. A direct consequence would be a higher average density of grid fields along the walls than in the center of the box.

### 5.4 Insertion of a wall

Our assumption on the origin of anchoring in the wall-removal experiment has the consequence that the reverse experimental paradigm (inserting a wall in a familiar open field) would not lead to the reverse results, i.e., grid fields would not reorganize into two incoherent grids. Instead, we predict that the insertion of the wall would introduce grid distortions close to the wall, maybe up to the formation of new fields due to the excitatory aspect of the wall.

## 6 Relationship between microscopic and macroscopic levels

Here we propose a macroscopic description, in the sense that we directly model firing patterns in physical space and not the underlying neurons. So far, with the notable exception of the study by Krupic et al. (2014) (see Discussion), all grid cell models were microscopic. It is thus crucial to bridge the gap between both levels of description. So we ask: (1) what is the outcome of microscopic models when looked at macroscopically? (‘bottom-up’ direction) And, conversely, (2) how can our macroscopic interactions be implemented in a neural network? (‘top-down’ direction).

### 6.1 From microscopic to macroscopic

#### 6.1.1 Attractor mechanisms

This class of continuous attractor network models (McNaughton et al. (2006); Fuhs and Touretzky (2006); Guanella et al. (2007); Burak and Fiete (2009); Pastoll et al. (2013); Couey et al. (2013); Widloski and Fiete (2014)) is based on the collective behavior of a large number of interconnected cells. A specific connectivity between neurons arranged on a 2-dimensional *‘neural sheet’* allows the emergence of activity patterns—called Turing patterns—that form a hexagonal grid on this neural sheet. These patterns have to be translated into physical space. The models therefore incorporate a mechanism that translates the activity on the neural sheet according to the motion of the animal in physical space (‘path integration’). This way, up to some noise correction, each cell has a hexagonal firing pattern in physical space.

We take the following reasoning: under the assumptions that periodic Turing patterns form in the neural space and that the path integrator is reliable enough (*i.e.* assuming the model actually works), we can directly convert the multi-cell / single-time description of the microscopic model into its single-cell / across-time equivalent at the macroscopic level. The reasoning is sketched hereafter.

##### In neural space

Monasson and Rosay (2014) have demonstrated formally that, in a 1-dimensional continuous attractor network with short-range excitation and global inhibition, the bump of activity that forms moves like a quasi-particle, thus validating a macroscopic description. More precisely, in the large *N* limit and in the presence of neural noise, across time the bump undergoes little deformation (scaling as 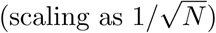) and its center diffuses (with a diffusion coefficient scaling as 1*/N*).

Though it would require a rigorous demonstration, we assume that at least the quasi-particle behavior remains true in the case of a 2-dimensional continuous attractor network with connectivity leading to grid patterns. Moreover, the large *N* limit is biologically plausible: from estimates of the number of neurons in the rodent entorhinal cortex (Merrill et al. (2001)) and the number of modules (Stensola et al. (2012)), one can deduce that *N* is of the order of 10^4^.

When forces (here, external inputs of velocity) are applied to the bump, Monasson and Rosay (2014) have observed that it keeps its quasi-particle aspect as long as the intensity of the input is not too large. So we expect it to hold also in the presence of path-integration forces moving a multiple-bump pattern.

##### In physical space

We argue that this description holds as soon as the model forms stable grids (see Supplementary Material). Moreover, there is a correspondence between the stable states of the neural network and the single-cell grid patterns, hence we can directly derive an approximate effective single-cell Hamiltonian at the macroscopic level. For instance, in the inhibition-based model by Couey et al. (2013), at the macroscopic level grid fields from the same grid cell repulse each other while their density is maintained by a chemical potential.

Here we are in the case where the effective forces between grid fields are a network effect, hence grid fields from another cell from the same module would be directly given by a translation of the grid fields from the previous grid cell.

In the merging setup, attractor models have not been simulated yet. We predict intuitively that after removing the wall a perfect grid would form that is an extension of the grid in the compartment where the rat is when the wall is removed. Therefore, some ingredient is missing to account for the observed merging of incoherent maps. More generally, an open question for attractor-based mechanisms is how specific grid patterns are selected that do not correspond to the ‘perfect and universal grid’ framework. This is not only necessary to explain the merging experiment, but also for error-correction, retrieval of the same grid maps upon re-entering the same environment, effects of walls and in general all cases of grid anchoring and distortion. Hardcastle et al. (2015) have shown that border cell-like excitatory inputs succeed in error-correction in an attractor model. Their assumption remains yet to be tested in grid-distortion paradigms.

#### 6.1.2 Adaptation mechanisms (and other competitive learning mechanisms)

Let us now turn to the class of models initiated by Kropff & Treves in 2008 (Kropff and Treves (2008); Si et al. (2012); Si and Treves (2013); Stella et al. (2013); Urdapilleta et al. (2015); Stella and Treves (2015)). Here, grid patterns are not a consequence of recurrent connectivity between a large number of cells, but come from a single-cell mechanism (adaptation), combined with a competition between place-modulated inputs. Adaptation tends to make the cell self-inhibit while the external inputs maintain its average level of firing and anchor the firing fields. Excitatory recurrent collaterals can be added on top of this system to account for module coherence (Figure 11a).

**Figure 11.**
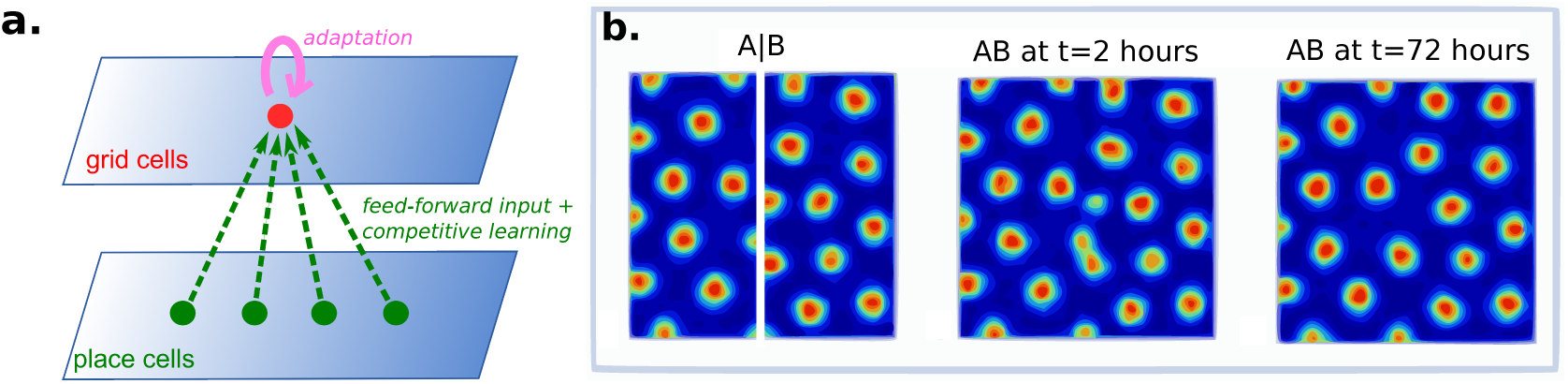
Adaptation-based models in the merging experiment. **a**: General principle: each grid cell tends to self-inhibit on a certain lapse of time, while receiving place-modulated inputs. When averaged over many trajectories, this results in an effective spatial kernel of repulsion between fields (Kropff and Treves (2008)). **b**: Example of a simulated grid cell of the model by Si et al. (2012) in the wall-removal setup. Left: Rate maps after 3 days of exploration in boxes with the separation wall. Center: Rate map of the same grid cell in the open field after 2 hours of exploration. Right: Rate map of the same grid cell in the open field after 72 hours of exploration

In the original paper from 2008, Kropff & Treves argued that their model could be simplified by considering that each cell’s firing rate minimizes a cost function:

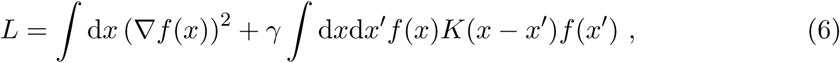

where *f* (*x*) is the cell’s firing rate at position *x* and *K*(*x – x’*) is an inhibitory kernel on a distance scale equal to the adaptation time constant times the average velocity of the animal. Moreover, *f* has to satisfy a constraint of constant average value.

Although this relationship was not formally derived in the paper—but the calculation was sketched by (Sprekeler (2008))—it basically boils down to a macroscopic effective version of the microscopic model. It is expressed in terms of firing rate instead of particle positions, but as soon as we know the solution is multipeaked (as shown in Kropff and Treves (2008)) we can replace the bumps by point particles with only a minor loss of information, and remove the irrelevant continuity term in the cost function. *K* corresponds to a repulsion between particles and the constant global activity constraint is the equivalent of our fixed number of particles. We therefore, here again, obtain an effective model with short-range repulsion and global excitation.

Nevertheless, this cost function description is valid in a limited domain only. First, it concerns only the simplest version of the model (without recurrent collaterals). Second, the presence of the walls is not taken into account while they do influence patterns by constraining the animal’s motion during learning. Finally, it only describes the asymptotic state of the firing pattern, and not its dynamics before equilibrium is reached.

Notably, in the case of manipulations of the environment’s shape, like wall removal, the adaptation model takes a very long time to stabilize because it is strongly anchored to the old configuration by the feed-forward inputs. We verified this aspect by simulating the model by Si et al. (2012) in the wall-removal setup (Figure 11b). Simulation details are provided in Supplementary Material.

The first thing we notice is that the evolution of the rate maps looks qualitatively very similar to the experimental ones: two nice-looking but incoherent grid maps before wall removal, then an evolution of the central fields towards more coherence. However, the durations do not match. Stabilization times vary from one simulation to the other, but are generally of the order of tens of ‘rat-equivalent hours’ (Figure 11b). In contrast, in experiments the merging process takes place within a few minutes (Wernle et al. (2018)). The inertia in the model due to the place-modulated input does not match the fast evolution of the grid patterns observed experimentally.

Recently, Weber and Sprekeler (2018a) introduced a model for grid cells based on an interaction of excitatory and inhibitory inputs through synaptic plasticity. Both types of inputs are place-modulated. Periodic patterns arise if the spatial tuning of inhibitory inputs is broader than that of excitatory inputs. In this case also, grid fields behave as if they repulse each other. Although different regarding its hypotheses at the cellular level, it is interesting to note that this model shares common points with the adaptation model. Its simulation in the wall-removal experiment also shows a local rearrangement of grid fields towards a more coherent pattern, although on a faster timescale than the adaptation model (around 5 hours).

#### 6.1.3 Oscillatory-interference models

In oscillatory-interference models, grid cells are modeled with membrane potential oscillations (theta rhythm) modulated by running speed in specific directions (Burgess et al. (2007); Giocomo et al. (2007); Blair et al. (2007); Hasselmo et al. (2007)). These models have the advantage to naturally explain phase precession, while other grid cell models do not.

At the macroscopic level, grids emerge as the intersection of three stripe-shape interference patterns, and would therefore be better described as waves rather than particles. The derivation of such a macroscopic equivalent is not straightforward, though.

It is unclear how module coherence, anchoring, grid distortions and response to wall removal could be accounted for by oscillatory-interference models without additional ingredients.

### 6.2 From macroscopic to microscopic

We now ask the reverse question: assuming that our macroscopic description is correct, what can we infer about the underlying microscopic system? The full derivation of a microscopic model is out of the scope of the present study, however, we can sketch the constraints that such a model would have to fulfill.

#### Existence of a description as interacting particles

If the dynamics of grid fields can be accurately described as interacting particles, this highlights that not only do grid cells have astonishingly simple firing correlates but also the relationships between them have very simple spatial expressions.

In a sense, this was already assumed in the framework of the perfect universal grid, where the relationship between grid fields from the same cell was a rigid triangular lattice; and the relationship between grid cells from the same module was given by a rigid offset. What we have shown here is that this view extends beyond perfect grids to distorted patterns. These distorted grids are not just noisy hexagons. Instead, they can be approximated by a system where forces only depend on spatial distances. This points to a remarkable conversion mechanism between distances and angles on the one hand, and neuronal dynamics on the other hand. This mechanism should be stable enough to cope with erratic trajectories, permissive enough to let other inputs distort the grids, and simple enough to be implemented in a neural network.

#### Grid symmetry

We have seen in Section 3 that some experimental grid patterns have extremely high levels of order, while others do not. In our macroscopic model, we could account for this by a quasi-crystalline state of a large number of interacting particles. But how is this implemented at the microscopic level? What kind of neural network produces all gridness scores ranging from −0.5 to 1.5, virtually their theoretical upper bound?

High gridness scores are an argument in favor of a many-cell effect: intuitively, crystalline orders can easily emerge when a lot of units are interacting together; while if there are only a few of them the order is much less robust to noise in the parameters. And indeed, the existing attractor-based models do produce high gridness scores (Bonnevie et al. (2013)).

Conversely, concerning the rest of the distribution of gridness scores, the roles are switched. So far only adaptation-based models have accounted for low gridness scores: from what has been reported, attractor-based models seem to produce *only* very high gridness scores. ^5^

These considerations guide us towards attractor-based models that incorporate localized inputs. Such inputs could suffice to distort the grids and explain the full range of observed gridness scores, the same way as local forces from the walls do lead to grid distortions in our macroscopic model. Such a *‘hybrid model’* remains to be implemented.

#### Repulsion between grid fields

The repulsive force between particles is the main ingredient behind grid pattern formation in our model. The fact that repulsion leads to hexagonal patterns is a quite general phenomenon in physics, see for instance Abrikosov vortices in superconductors materials (Abrikosov et al. (2012)). Many different microscopic mechanisms could lead to an effective repulsion between fields: recurrent connectivity in a neural sheet, adaptation, inhibition from other non-grid place-modulated inputs, etc. On this aspect, the macroscopic approach does not put much constraint on the microscopic level.

#### Inter-cell coupling

We have shown in Section 4.2 that grid cells of the same module can be described by introducing a weak coupling between grid fields of different cells (including an offset). We also have shown that, if cells form their patterns independently (simplified as perfect hexagons) and those patterns repulse each others, then the orientations align and the phase are evenly dispersed. This again points towards a hybrid-model: either a single-cell pattern formation model on top of which coupling between cells is added, or an attractor network model on top of which local inputs are added. Further quantitative measures of grid distortion and module coherence would help to distinguish between them.

#### Interaction with the walls

Excitatory border cells could implement the orthogonal attraction we used. However, that attraction is not the only possibility. We expect upcoming experiments to shed more light on what walls do to grids, notably by performing local measures.

## Discussion

### Summary of results

A growing body of experimental evidence shows that the idealized view of perfect and universal grids has to be abandoned. For the same reason, the gridness score measure—that was based on this idealization—is not the optimal way to characterize grid patterns that are prone to distortions, inhomogeneities and absence of long-range order. Taking inspiration from the rich field of Physics of two-dimensional systems and colloids, we have proposed both an alternative measure for grid patterns that is simple and local and a macroscopic model for their formation. We have shown that this model, although very simple, reproduces most of the observed features of grid patterns.

One of our main results is that both local hexagonal order and long-range breaking are necessarily a large-*N* effect. Simulating a large number of grid fields creates grid patterns that share many aspects with experimental recordings.

We then turned to the multiple-cell case. Remaining agnostic about whether or not the module’s recurrent connectivity is already contained in the effective interaction between grid fields, we have shown that a relatively small force is enough to align two patterns. We have also seen that in this model, hexagonal symmetry and module coherence are two independent features that can be separated, a prediction that could be tested experimentally. We then tested the assumption that inhibition between grids would lead to both orientation alignment and phase dispersion in a simplified setup by simulating repelling hexagonal patterns. We did observe the expected phenomena. The interaction with pattern formation remains to be tested in simulations combining both large numbers of particles and large numbers of grid cells.

The force that walls exert on the grid pattern is not fully understood, both at the experimental level and in our effective model. We have preliminary indications that attractive walls lead to patterns that qualitatively resemble experimental recordings, with phenomena such as bending of grid axes and barrel distortion. Hopefully, future experiments will clarify those phenomena.

In the case of the wall-removal setup, we drastically simplified our assumptions and showed that even a minimal implementation captures the main features of the experimental observations. Notably, it is not necessary to assume a force coming from the walls to account for Wernle *et al* ‘s results.

### Significance

The use of effective, macroscopic models is not common in neuroscience. The reason is probably that computational neuroscience aims at explaining brain observables from the neurons themselves, and therefore places itself at the level of neurons. Our point here is that both approaches are complementary. A macroscopic model describes rather than explains, but an accurate description is a prerequisite to any attempt of an explanation. Note that almost all characterizations of grid cells have been done at a macroscopic level: ratemaps, autocorrelograms, gridness scores, shearing, grid field displacement, etc. Grid cells even take their name from their macroscopic features. Here we refined and systematized a natural approach to describe grid patterns.

This way, we have sketched the correspondence between microscopic and macroscopic levels of description. We have seen that the microscopic mechanisms previously proposed are either incomplete (for the attractor-based and oscillation-based models) or too slow (for the adaptation-based). A hybrid model between attractor mechanisms and single-cell effects seems a good candidate. We argue that a good description at the macroscopic level is useful to guide the design of a microscopic model.

Moreover, working on the macroscopic level is fast to simulate, requires few parameters and allows to make testable experimental predictions. We provided some of them here, but extensions like curved spaces, higher dimensions, fancy geometries, etc., would be straightforward to implement.

### Criticisms

As any simplification, our model misses some features. For instance, the reduction of bumps of activity to point particles cannot account for different peak firing rate that, as Dunn et al. (2017) recently showed, could be crucial in terms of coding. This aspect could nevertheless easily be added to the model by assigning different potential weights to the different particles. We also lose the information on timescales of the dynamics, as illustrated by our simulations of the wall-removal experiment in the adaptation model. We do not account for the merging of two grid fields observed in some cases by Wernle *et al*: this could be corrected by adding a short-range attractive force between grid fields, a creation / deletion process and some chemical potential controlling the total number of particles. We chose to stick to simplicity by discarding this aspect.

### Comparison with the model by Krupic et al (2014)

Krupic *et al* (2014) also suggested a model of interacting grid fields. They assume a force between grid fields that is derived from a Lennard-Jones potential (i.e., long-range attraction, short-range repulsion). The walls exert a repulsion on the fields, orthogonal to the walls with an exponential decay. Even if similar in spirit, the two models have a crucial difference: while we see pattern formation as a self-organized, emerging feature of a large number of grid fields, in the model by Krupic *et al* it is the result of fine-tuning on a small number of particles: the range of the Lennard-Jones potential, of the repulsion from the walls and the number of particles has to be carefully chosen *ad hoc* to produce hexagons. As a result, either only very good grids or very poor grids are observed (see their Fig.2), but not the whole distribution. At the microscopic level, Krupic *et al* sketched an implementation based on repulsion between CA1 place cells and inhibitory border cells—in contrast to our own conclusions.

### Extensions

The approach of a macroscopic model opens the way for several lines of future research.

The first concerns data analysis. The *Ψ* score introduced above is a promising way to characterize grid patterns, but it still remains to be extensively tested on experimental data. First encouraging results have just been reported by Weber and Sprekeler (2018b). It would be interesting to fit our model’s parameters and to compare the best fit with other versions of the model (e.g. other shapes of the interactions). From this, two paths could be taken: making more precise predictions on the outcome of future experiments, and refining our inference on the underlying microscopic system by a better discrimination between the candidate mechanisms.

Our effective approach could also be extended to other spatially tuned cell types, or in studying the still mysterious relationship between grid modules. More generally, the spirit of an effective model is not limited to the hippocampal formation and could be fruitful in the study of many other brain areas.

Finally, since we still want to understand what grid cells—rather than grid fields— do, developing a complete microscopic model is the ultimate step. We have seen that the question is not so much how to get good grids, but how to get both excellent and poor grids. We stressed that a hybrid mechanism combining ingredients from the different existing models is a good candidate. We hope that the hindsight provided by the macroscopic point of view will be of help when diving in the intricate complexities of the microscopic world.

## Supporting information

## Acknowledgements

We are grateful to Alessandro Treves, Rèmi Monasson, Giuseppe D’Adamo, Thomas Gueudrè and Henning Sprekeler for their remarks on the model and its relationship with Physics. We thank Tanja Wernle for extensive discussion on the merging experiment. S.R. would like to thank the GRIDMAP project for financial support and the Abdus Salam International Centre for Theoretical Physics in Trieste for hospitality in the conclusive phase of this work. Many thanks also to John Nicholls. The idea of grid alignment via recurrent inhibitory connections was developed in discussions between Henning Sprekeler and Simon Weber.

## Footnotes

1 Grid patterns with non homogeneous orientation have been shown by Stensola et al. (2015) but, in the absence of a local measure, could not be quantified.

2 Some slight differences, either of spacing or orientation, have been reported, but we neglect them for the present discussion.

3 A question not answered by this experiment is whether this position is fixed with respect to the walls or with respect to something else, e.g., distal cues.

4 Near the walls some grid distortion could come into play as we have shown, but let us assume here that the box is big enough so that we can neglect such edge effects.

5 Maybe noise could account for lower gridness scores in adaptation models, but to our knowledge this possibility has not been investigated yet.

