## Supplementary material for "Modeling grid fields instead of modeling grid cells"

Sophie Rosay[1], Simon N. Weber[2], and Marcello Mulas[3]

[1]Scuola Internazionale Superiore di Studi Avanzati, Trieste (Italy)

[2]Technische Universität, Berlin (Germany)
[3]Technische Universität, Munich (Germany)

November 28, 2018

### 1 Simulation methods

#### 1.1 Algorithm

The simulations are run as follows: at each time step, for each particle, compute the forces exerted on it and update its velocity by integrating the Langevin equation

$$m\frac{\mathrm{d}\vec{v}_i}{\mathrm{d}t} = \sum_j \vec{F}_{j\to i} + \sum_k \vec{F}^{\mathrm{wall}}_{k\to i} - \alpha\vec{v} + \vec{\xi}(t) \ . \tag{1}$$

Asymptotically, after a few hundreds of steps necessary to stabilize, we get the final configuration (positions of the point particles). From it we can extrapolate the rate map, autocorrelogram and gridness score as detailed hereafter.

To obtain **rate maps** from field positions $\{(x_i, y_i)\}_{i=1\dots N}$, we just sum Gaussian bumps centered on these positions:

$$r(x,y) = \sum_{i=1}^{N} e^{-\frac{(x-x_i)^2+(y-y_i)^2}{2\sigma^2}} \tag{2}$$

where the bump width $\sigma$ is taken as $L_{\mathrm{rep}}/5$, leading to a realistic ratio between bump width and grid spacing.

The **autocorrelograms** are then computed from the rate maps by following the standard procedure, see e.g. **?**:

$$AC(\tau_x,\tau_y) = \frac{\Big\langle r(x,y)\cdot r(x-\tau_x,y-\tau_y)\Big\rangle_{x,y} - \Big\langle r(x,y)\Big\rangle_{x,y}\Big\langle r(x-\tau_x,y-\tau_y)\Big\rangle_{x,y}}{\sqrt{\Big\langle (r(x,y)-\langle r\rangle)^2\Big\rangle_{x,y}\Big\langle (r(x-\tau_x,y-\tau_y)-\langle r\rangle)^2\Big\rangle_{x,y}}} \quad (3)$$

where the averages are taken over the overlap area between the original map and the shifted one. The autocorrelogram is defined only for translations $\tau$ for which the overlap is greater than 20 bins.

The computation of the **gridness score** from the autocorrelogram takes the following steps:

- remove an inner disk around the central peak of the autocorrelogram, of radius $R_{\text{in}}$ defined from the most central region of value greater than 0.2.
- remove an outer disk of increasing radius
- at each step of the radius length $R_{\text{out}}$,
  - Rotate the autocorrelogram ring thus obtained by an angle $\psi = 30, 60, 90, 120$ and 150 degrees successively. - At each rotation $\psi$, compute the correlation $\text{Corr}(\psi, R_{\text{out}})$ between the original ring and the rotated one.
  - Compute

  $$G(R_{\text{out}}) \equiv \min\Big(\text{Corr}(60,R_{\text{out}}),\text{Corr}(120,R_{\text{out}})\Big) - \max\Big(\text{Corr}(30,R_{\text{out}}),\text{Corr}(90,R_{\text{out}}),\text{Corr}(150,R_{\text{out}})\Big) \quad (4)$$

- the gridness score is equal the maximal value of G over the range of the external radius $R_{\text{out}}$.

### 1.2 Parameters and shape of the interactions

In the simulations reported above, we use a Gaussian kernel for the forces. Therefore the *repulsive* forces between particles take the form:

$$\vec{F}_{j\to i} = R_{\text{rep}} \exp\left(-\frac{d_{ij}^2}{2L_{\text{rep}}^2}\right)\vec{u}_{ij}\ , \quad (5)$$

where $d_{ij}$ is the distance between particles $i$ and $j$ and $\vec{u}_{ij}$ is a unitary vector from $j$ to $i$. Similarly in the case of the *attractive* coupling between two cells from the same module:

$$\vec{F}_{j\to i}^{\text{coupl}} = -R_{\text{coupl}} \exp\left(-\frac{d_{ij'}^2}{2L_{\text{coupl}}^2}\right)\vec{u}_{ij'}\ , \quad (6)$$

where $j'$ indicates the position of particle $j$ translated by the assumed grid offset (taken at random in simulations).

The forces exerted by the walls on the particles write:

$$\vec{F}^{\text{wall}}_{k\to i} = -R_{\text{wall}} \exp\left(-\frac{d_{ik}^2}{2L_{\text{wall}}^2}\right) \vec{u}_{ik} \ , \tag{7}$$

where $d_{ik}$ is the distance between particle $i$ and the wall $k$ and $\vec{u}_{ik}$ is a unitary vector orthogonal to wall $k$ ($R_{\text{wall}}$ is *positive* for an *attractive* force).

We used the following parameter values:

| Figure | $R_{\text{rep}}$ | $L_{\text{rep}}$ | $R_{\text{coupl}}$ | $L_{\text{coupl}}$ | $R_{\text{wall}}$ | $L_{\text{wall}}$ | $\alpha$ | $\beta$ | $L$ | $N$ |
|---|---|---|---|---|---|---|---|---|---|---|
| Fig. 4a | 1 | 0.2 | / | / | 1 | 0.2 | 10 | 100 | 1 | 25 |
| Fig. 4b | 1 | 0.5 | / | / | / | / | 10 | 100 | 10 | 1000 |
| Fig. 5 | 1 | 0.5 | / | / | / | / | 10 | 100 | 10 | 1000 |
| Fig. 6 | 1 | 0.5 | varies | 0.3 | / | / | 10 | 100 | 10 | 1000 |
| Fig. 8a | 1 | 0.5 | / | / | 0.2 | 0.5 | 10 | 100 | 10 | 1000 (of which 43 inside) |
| Fig. 8b | 1 | 0.7 | / | / | 0.2 | 0.7 | 10 | 100 | 10 | 1000 (of which 20 inside) |
| Fig. 9 & 10 | 1 | 0.4 | / | / | / | / | 10 | 100 | 2 | 28 (+ fields outside) |

Table 1 Parameter values used in the simulations reported throughout the present study

The formation of grid pattern is robust on a wide region of these parameters. The increase of temperature (*i.e.* decrease of $\beta$) and the decrease of the density of fields lead to a fluid phase. Increasing the density beyond the density of soft spheres optimal packing also destroys the order. We have also tested other forms for the potential $U_{ij}$ as a function of the distance between particles: a power-law decay, a Gaussian decay and a Mexican-hat shape (results not shown here). In each case we found parameter values accounting for grid formation.

### 1.3 Measure of coherence between two patterns

In order to quantify how coherent two patterns are, i.e. how close one is to be a translation of the other, we define a measure of their 'incoherence' in the following way:

- translate one pattern relatively to the other in all possible direction within the grid scaling;
- for each translation $T$, for each translated grid field $i_1$ of pattern 1, find the closest grid field $i_2$ of pattern 2;
- sum the distances between all the thus found pairs of fields:

$$S(T) = \sum_{i_1} \min_{i_2}(\text{dist}(T(i_1), i_2)) \ , \tag{8}$$

where $T(i_1)$ denotes the translation of field $i_1$.

- find the translation $\hat{T}$ that minimizes $S(T)$.
- the module 'incoherence' $I_{\mathrm{mod}}$ is defined as the mean distance between pairs of fields after this translation, normalized by the grid spacing:

$$I_{\mathrm{mod}} \equiv \frac{1}{NL} \sum_{i_1} \mathrm{dist}(\hat{T}(i_1), i_2) = \frac{1}{NL} S(\hat{T}) \ . \tag{9}$$

### 1.4 Simulations of interacting perfect hexagons with repulsion

We now describe how we obtained the phases and orientations that minimize overlap between grid fields in Fig. 7 in the main text. We create $N$ perfect hexagonal grids of spacing 1 with 91 fields each (Fig. 1 shows a smaller illustration with 19 fields). The first grid is fixed, with phase $(0,0)$ and orientation 0. We define an energy function that quantifies the overlap of all $N$ grids within a circle of radius 3 that is centered at the central node of grid 1 (gray circle shown in main text Fig. 7a). If not for this constraint, shifting two grids by a complete spacing would reduce the overlap of all fields at the boundaries of the grids, which is not a minimum that we are interested in (Fig. 1). To obtain the overlap, we compute the pairwise distances between all fields $i$ of grid $k$ with all fields $j$ of grid $l$ for all $N$ grids, including only fields that lie within the circle. We assume grid fields to be Gaussians, and since the overlap between two Gaussians is again a Gaussian, we define the energy as

$$E = \sum_{k<l}^{N} \sum_{i \in f(k)} \sum_{j \in f(l)} \exp\left\{ -\|\mathbf{x}_{k,i} - \mathbf{x}_{l,j}\|^2 / 2\sigma^2 \right\} ,$$

where $\|\cdot\|$ denotes the euclidean length and $\mathbf{x}_{k,i}$ is the center location of the $i$-th field of grid $k$. The sum over $k < l$ indicates that we do not compare grids twice and do not consider the overlap between fields of the same grid. The set of indices of field centers of grid $k$ that lie inside the circle of radius 3 is denoted by $f(k)$. Moreover, $\sigma$ determines the width of individual fields. Choosing $\sigma$ too large makes shifts effectless, whereas choosing $\sigma$ too small leads to many optimal solutions in which grid fields do not overlap at all. We set $\sigma = 0.2$, which leads to meaningful results for all $N$ under consideration. Note that $E$ is a function of the phases and orientations of the $N-1$ non-fixed grids. We minimize $E$ using a standard optimizer (`scipy.optimize.minimize` from the *SciPy* package for *Python*) starting from a random initialization of phases. In one set of simulations, we fix all orientations at 0 degrees and minimize the energy with respect to $N-1$ phases. In a second set of simulations, we minimize the energy with respect to $N-1$ phases *and* orientations, but initialize all orientations at 0 degrees. In a third set of simulations, we minimize the energy with respect to $N-1$ phases *and* orientations and initialize all values at random. Grids with identical orientations consistently result in lower energies (main text Fig. 7a,b). Minimizing with respect to phases *and* orientation, without initializing at 0 degrees, typically gets stuck in a local minimum without identical orientation (main text Fig. 7b). This analysis shows that an arrangement of identical orientations is a deep local minimum. Whether it is the *global* minimum remains to be shown.

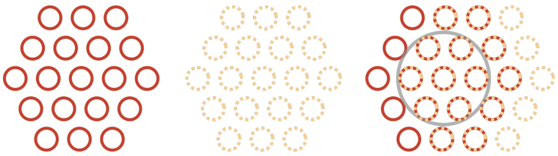

Figure 1 Illustration of why we consider only the overlap within a circle in the minimization problem. Two perfect grids (left and center) of 19 fields each. If the two grids are phase shifted by one grid spacing with respect to each other, their overlap is reduced; only 14 field overlap (right). Within the gray circle, a phase shift of one grid spacing does not reduce the overlap

#### 1.5 Simulations of the adaptation model in the wall-removal setup

We investigated the merging process of spatial grid patterns generated in separated compartments by simulating populations of 50 conjunctive cells with the model by **?**.

This model is capable of reproducing the emergence of conjunctive grid-by-head-direction cells by exploiting firing rate adaptation. Grid-like activity of single neurons emerges with the interplay of plastic feed-forward projections from a population of place cells and internal recurrent connections.

In addition to the neural activity we simulated a rat randomly exploring a $2\times2$m square environment with a removable wall in the middle. The wall separated two identical compartments named ‘A’ (on the left) and ‘B’ (on the right). In sequence, the rat first explored A for 1.5 days, then B for 1.5 days and, lastly, after removing the separation wall, it freely explored the open space ‘AB’ for 3 days. With a time step equal to 10 ms, 3 days of simulated time corresponded to 25920000 simulation iterations.

Main text’s Fig. 11 shows rate maps from one example simulation. Supplementary Figure 2 shows the average cross-correlation between the maps generated before and after the removal of the separation wall as a function of the distance from the wall. Lower values of cross-correlation indicate greater levels of grid reorganization. As expected, the reorganization occurred more at the interface between the two compartments (at distance equal to 0) rather than far from it (at distance equal to -1 or 1). The grid merging process was entirely due to the plasticity of the feed-forward projections. In fact, it occurred even in absence of collateral connections ($\rho = 0$). However, the stronger the collaterals, the more rigid the spatial maps. As Figure 2 shows, an increase in the strength of collaterals $\rho$ resulted in an increase of the average cross-correlation between maps.

### 2 Attractor mechanism: derivation of the corresponding macroscopic model

Here we detail the reasoning sketched in Section 6.1.1., providing an approximate macroscopic description of attractor-based models.

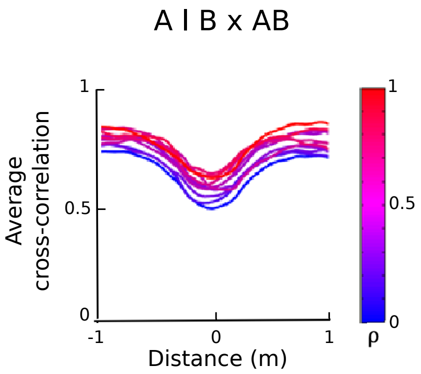


Figure 2 Simulations of the adaptation model by **?** in the wall-removal setup: average cross-correlation as a function of the distance from the separation wall of 50 spatial maps independently obtained in the compartments A and B ('A—B') and in the open space 'AB' for different values of the collateral strength $\rho$

### 2.1 Can such a model be described in terms of interacting quasi-particles?

In neural space, we know from **?** that, in a continuous attractor network with no external forces, the bump of activity diffuses with little deformation, hence behaving as a colloidal particle. So, if the animal is at a constant position $x_0$—no path integration input—let us write (the hat indicates quantities in neural space):

$$\hat{\rho}_{x_0}(i,t) \approx \hat{\rho}_{x_0}(i - \Delta i(t), t_0) \tag{10}$$

where $\hat{\rho}(i,t)$ is the shape of the activity bump at time $t$ in the neural space parametered by neuron index $i$. Its deformations across time have been neglected; it only undergoes a translation of its center, obeying a pure diffusion equation.

In the case of several bumps (that would be the case of a grid cell model), things are a bit more complicated because other modes would have to be taken into account, but we expect that in first approximation the particle-like description would still hold.

Let us assume that (a) the path-integrator is accurate and has no noise and (b) the typical time of diffusion of the Turing pattern $\tau_{\text{diff}} = 1/D$ is much longer than the time the animal takes to explore the whole environment $\tau_{\text{mov}} \sim L/v$. Because we neglect the deformations of the bump, the quasi-particle diffusion in neural space results in a quasi-particle diffusion in physical space. Ideed in physical space we can write:

$$\rho_{i_1}(x_1) = \hat{\rho}_{x_1}(i_1) = f(x_1, i_1) \tag{11}$$

The path integration mechanism is written

$$f(x_1 + \Delta x, i_1) = f(x_1, i_1 - \Delta i) \ , \tag{12}$$

$$\Delta i = g_{\text{PI}} \cdot \Delta x \ , \tag{13}$$

where $g_{\mathrm{PI}}$ is the gain of the path integrator.

Therefore, when in neural space the Turing pattern diffuses say of a quantity $\Delta i$, in physical space it converts into

$$\rho'_{i_1}(x) = \hat{\rho}'_x(i_1) = \rho_{i1}(x + \Delta x) \tag{14}$$

In other terms, because of the linearity of the path integration mechanism, a diffusion of the Turing pattern in neural space leads to a diffusion of the grid pattern in physical space.

The two assumptions we used to establish this result are the accuracy of the path integrator and the slow diffusion timescale as compared to the animal's motion:

- The possibility of a noisy path integrator has actually already been the topic of previous studies (**??**). Above a certain level of noise the attractor network does not produce good grids anymore, hence it is also necessary for the model to work to assume that the path integrator is not too noisy.
- What happens if the Turing pattern diffuses faster than the animal's motion? Here again, a consequence would be the loss of the grid pattern in physical space. Therefore, we can reasonably consider that our quasi-particle reasoning is valid in the domain where the attractor network actually works.

### 2.2 What are then the effective forces on particles?

**Grid fields from one grid cell** Let us first consider one given grid cell.

Like before, let us assume that the path integrator is accurate and that the diffusion in neural space is slower than the exploration time. Moreover, Turing patterns are stable states and therefore the observed patterns of activity in neural space only differ by a translation. Under these assumptions, we are in the case of a pure reproduction of the same pattern in neural space (single time, many cells) in physical space (across time, single cell). In other words, the pattern of grid fields of the grid cell we are looking at is exactly given by the pattern of activity in neural space. To fix ideas, let us for instance consider the model by **?**.

In neural space, if we neglect the velocity input, we can estimate that the activity is driven by the Hamiltonian:

$$\hat{H} = -\sum_{i<j} J_{ij} f_i f_j - I \sum_i f_i \ , \tag{15}$$

where $f_i$ is the firing rate of neuron $i$, $J_{ij}$ is the synaptic coupling between neurons $i$ and $j$ (uniformly inhibitory on a disk of radius $R_0$) and $I$ is the global excitation coming from the hippocampus.

From the argument above, we can say that in first approximation, and in the regime of reliable path integration, the grid cell we observe has patterns of activity governed by the Hamiltonian

$$H = -\int \mathrm{d}x\mathrm{d}y J(x,y) f(x) f(y) - I \int \mathrm{d}x f(x), \tag{16}$$

where $J(x, y)$ is equal to a negative constant on a disk of radius $R_0/g_{\mathrm{PI}}$.

So we can see the Turing pattern in physical space as the result of local inhibition between the bumps. The combined result of the network attractor dynamics and the path integration mechanism is that the grid fields from one cell exert an effective repulsion on their neighbors and undergo a global excitation: the effective interactions between fields in physical space mirror the connectivity pattern in neural space. This comes from the fact that the accurate path integration hypothesis allows us to directly translate [neural-space / multiple-cell / single-time activity] into [physical-space / single-cell / across-time activity].

If we see those activity bumps as quasi-particles, the model (eq.16) has the same shape as ours (eq. 1 in Main Text), with step-shaped inhibition, no forces from the wall, and a chemical potential term.

**Grid fields from other grid cells from the same module** Our above conversion from neural space to physical space assumes that the single-cell effective model contains all the network's dynamic. And indeed, given the firing pattern of a grid cell, the microscopic model strongly constrains the firing of any other cell from the same module. The only way they can depart from being a mere translation of the other cell's pattern is through noise (in neural firing or in path integration) that would be averaged out if considered on long time scales. Therefore, given the firing pattern of the single cell, all the other cells from the same module 'see' a deep 'egg box' potential given by the single cell's pattern.

**Grid alignment and anchoring** As pointed out above, the model by **?** does not contain inputs localized in space to ensure the resetting of the path integrator and the anchoring of the grids.

A possible mechanism for grid alignment would be through an input from border cells, as proposed by **?**. Another straightforward candidate for grid anchoring is the place cell system from the hippocampus, in a way that is still to be demonstrated.

Hence the questions: 'How do grid cells retrieve the same offset and orientation each time the animal enters the environment again? Is it related with grid distortions?' leave for future studies an interesting point to address.
